# CBF-1 promotes the establishment and maintenance of HIV latency by recruiting Polycomb repressive complexes, PRC1 and PRC2, at HIV LTR

**DOI:** 10.1101/174607

**Authors:** Sonia Zicari, Kalamo Farley, Lin Sun, Liam Spurr, Andrea Dragon, Michael Bukrinsky, Gary Simon, Ashok Chauhan, Mudit Tyagi

**Affiliations:** Division of Infectious Diseases, Department of Medicine, George Washington University, Washington, DC; Section of Intercellular Interaction, Eunice Kennedy Shriver National Institute of Child Health and Human Development, National Institutes of Health, Bethesda, MD, USA; Department of Microbiology,Immunology and Tropical Medicine, George Washington University, Washington, DC 20037

**Keywords:** HIV-1, latency, epigenetics, CBF-1, PRC1, PRC2, chromatin, transcription

## Abstract

The C-promoter binding factor-1 (CBF-1) is a potent and specific inhibitor of the HIV-1 LTR promoter. Here we demonstrate that the knockdown of endogenous CBF-1 in latently infected primary CD4+ T cells, using specific small hairpin RNAs (shRNA), resulted in the reactivation of latent HIV proviruses. Chromatin immunoprecipitation (ChIP) assays using latently infected primary T cells and Jurkat T-cell lines demonstrated that CBF-1 induces the establishment and maintenance of HIV latency by recruiting Polycomb Group (PcG/PRC) corepressor complexes or Polycomb repressive complexes 1 and 2 (PRC1 and PRC2). Knockdown of CBF-1 resulted in the dissociation of PRCs corepressor complexes enhancing the recruitment of RNA polymerase II (RNAP II) at HIV LTR. Knockdown of certain components of PRC1 and PRC2 also led to the reactivation of latent proviruses. Similarly, treatment of latently infected primary CD4+ T cells with the EZH2 inhibitor, 3-deazaneplanocin A (DZNep), led to their reactivation.

## Introduction

The anti-HIV therapy, ART has been highly successful in controlling Human Immunodeficiency Virus (HIV) replication and maintaining the level of HIV below the limit of detection (Gulick et al., 1997; Perelson et al., 1997). However, interruption of ART, even after decades of successful anti-HIV therapy, results in rapid and robust viral rebound (Chun et al., 1999; Finzi et al., 1999; Wong et al., 1997). The failure of ART to eradicate HIV is due to the creation of stable reservoirs of latently infected cells harboring slowly or non-replicating viruses. The majority of latent proviruses resides in resting memory CD4+ T cells, which provide a stable pool of latently infected cells with half-life roughly around 44 months (Brennan et al., 2009; Joos et al., 2008; Siliciano et al., 2003). These latent reservoirs are frequently replenished due to both homeostatic proliferation of latently infected cells and ectopic reactivation of latent proviruses followed by new rounds of infection, presumably owing to locally sub-optimal ART concentrations (Chomont et al., 2009; Chun et al., 2005; Hosmane et al., 2017; Tyagi and Bukrinsky, 2012). It is now well established that ART alone cannot eradicate latently infected cells, since intensification of ART was also found ineffective in reducing latent reservoir (Chun et al., 2005; Dinoso et al., 2009). Hence, developing therapeutic interventions with a focus on HIV eradication will require precise definition of the molecular mechanisms responsible for both the establishment and maintenance of HIV latency in order to either reactivate latent proviruses so that they can be destroyed, or fossilize them forever like Human Endogenous Retroviruses (HERVs).

As a retrovirus, the replication of HIV depends on efficient transcription. HIV transcription primarily relies on the availability of the host cell transcription machinery along with HIV own master transactivator protein Tat. Hence, flaws in proviral transcription appear to be the major factor contributing to HIV latency. Numerous factors and multiple mechanisms are known to impair HIV transcription and thus shown to promote HIV latency (Mbonye and Karn, 2014; Siliciano and Greene, 2011; Tyagi and Bukrinsky, 2012; Tyagi and Romerio, 2011). Notably, the type of epigenetic modifications and the resultant state of chromatin structures at the integrated HIV provirus provides critical signals that regulate transcription during both productive and latent HIV infections (Hakre et al., 2011; Karn, 2011; Margolis, 2010; Tyagi and Bukrinsky, 2012). The role of repressive epigenetic modifications in supporting HIV latency is quite evident by the fact that their removal or inhibition leads to the reactivation of latent proviruses (Choudhary and Margolis, 2011; Hakre et al., 2011; Margolis, 2011).

We previously described the important role of CBF-1, a CSL (CBF-1, SuH and Lag-1) type transcription factor, in restricting HIV transcription during HIV latency. CBF-1 is a key effector of Notch signaling pathways, which plays critical role in several developmental processes (Lai, 2002). CBF-1 restricts the expression of several cellular genes that carry appropriate DNA-binding sites for CBF-1 by recruiting histone deacetylases (HDACs) containing corepressor complexes (Borggrefe and Oswald, 2009; Ehebauer et al., 2006; Tyagi and Karn, 2007). By performing experiments in both transformed and primary CD4+ T cells we have established the role of CBF-1 as a potent repressor of HIV transcription (Tyagi and Karn, 2007; Tyagi et al., 2010). We have demonstrated that CBF-1, after binding to specific sites in HIV LTR, recruits corepressor complexes containing HDACs (HDAC1 and HDAC3). HDACs subsequently deacetylate the core histones and facilitate the establishment of transcriptionally repressive heterochromatin structures at HIV LTR. The closed/compact heterochromatin structures restrict the flow of transcriptional machinery at LTR promoter and thus hamper HIV transcription and promote HIV latency (Tyagi and Karn, 2007; Tyagi et al., 2010).

The recent literature suggests that CBF-1 restricts cellular gene expression not only through histone deacetylation, but also by inducing numerous other repressive epigenetic modifications, including trimethylation of histone H3 at positions lysine 9 (H3K9me3) or lysine 27 (H3K27me3) (Liefke et al., 2010; Martinez et al., 2009; Merdes and Paro, 2009). The presence of H3K9me3 and H3K27me3 at LTR and their role in establishing heterochromatin during HIV latency have already been demonstrated both by us and others (du Chene et al., 2007; Imai et al., 2010; Marban et al., 2007; Pearson et al., 2008; Tyagi and Karn, 2007). Notably, we have validated their physiological significance by showing the role of these repressive epigenetic modifications in establishing HIV latency in primary CD4+ T cells (Tyagi et al., 2010). Formation of H3K9me3 is primarily catalyzed by two kinases, namely SUV39H1 and G9A (Kouzarides, 2007). The methylation of histone H3 at position 27 (H3K27me3) is mainly catalyzed by EZH2 and occasionally by EZH1 (Cao et al., 2002; Kouzarides, 2002; Margueron et al., 2008; Shen et al., 2008). EZH2 and EZH1 are the main catalytic components of PRC2 complex (Cao and Zhang, 2004; Margueron and Reinberg, 2011). SUV39H1 and G9A frequently interact with different components of PRC1 complex (Li et al., 2010; Sewalt et al., 2002). PRC1 complex primarily inhibits transcriptional elongation via monoubiquitination of histone H2A, but it is also involved in several other epigenetic transactions along with the PRC2 complex via different interactions among their subunits (Cao et al., 2005; Lavigne et al., 2004; Margueron and Reinberg, 2011). PRCs play an important role in both inducing and maintaining the silencing of several cellular genes. PRCs restrict cellular gene expression by simultaneously inducing various types of repressive epigenetic modifications involving both histones and DNA, as PRCs carry multiple chromatin modifying enzymes (Beisel and Paro, 2011; Enderle et al., 2011; Kuzmichev et al., 2002; Margueron and Reinberg, 2011; van der Vlag and Otte, 1999; Vire et al., 2006). Consequently, PRCs-mediated epigenetic modifications regulate not only the transient gene silencing but also long-term silencing of the genes, such as of *Hox* genes and X-chromosome inactivation (Hernandez-Munoz et al., 2005; Okamoto et al., 2004; Plath et al., 2003; Silva et al., 2003).

In this study, we show that CBF-1 is the factor that promotes the recruitment of PRCs at HIV LTR. Recently, the role of PRCs during both the establishment and maintenance phases of HIV latency has been confirmed, and the presence of H3K27me3 at HIV LTR has been documented (Friedman et al., 2011; Matsuda et al., 2015; Nguyen et al., 2017; Tripathy et al., 2015). We further established the physiological significance, by showing the role of H3K27me3 during HIV latency in primary CD4+ T cells (Tyagi et al., 2010). However, the identity of the factor that recruits PRCs at HIV LTR was obscure. Remarkably, most of the enzymes that catalyze epigenetic modifications are not able to bind directly to the DNA and thus need to be recruited to DNA templates by various DNA binding proteins. Proteins such as CBF-1, YY1/LSF1, P50 homodimer, AP4, CTIP2, and thyroid hormone receptor recruit chromatin modifying enzymes in the form of multiprotein corepressor complexes to HIV LTR (Coull et al., 2000; Hsia and Shi, 2002; Imai and Okamoto, 2006; Marban et al., 2007; Tyagi and Karn, 2007; Williams et al., 2006). Using latently infected primary CD4+ T cells, we found that CBF-1 is the protein which recruits both PRC1 and PRC2 at HIV LTR. We confirmed that, by recruiting PRCs at HIV LTR, CBF-1 supports both the establishment and maintenance of HIV latency. Furthermore, we have validated the direct role of PRCs in HIV latency, as their knockdown results in the reactivation of latent HIV. Notably, very recently Karn group has demonstrated the role of JARID2 in recruiting PRC2 at HIV LTR (Nguyen et al., 2017).

## Materials and methods

### Cell culture, Cell lines and Chemicals

The CD4^+^ cells were isolated either from tonsils obtained from routine tonsillectomy or from peripheral blood mononuclear cells (PBMCs) of healthy donors, using Ficoll-Hypaque (GE Healthcare, USA) gradient centrifugation. CD4^+^ T cells were purified by negative selection method using a MACS kit (Miltenyi Biotechnology, Auburn, CA). CD4^+^ primary T cells and H80 cells were maintained in RPMI 1640 medium supplemented with 10% fetal calf serum and 25 mM HEPES. CD4^+^ primary T cells were supplemented with recombinant human IL-2 (20 U/ml) (R&D Systems, Inc.). 293T cells were grown in Dulbecco’s modified Eagle’s medium (DMEM) supplemented with 10% fetal bovine serum. T-cell lines CEM and Jurkat were maintained in RPMI 1640 medium supplemented with 10% fetal bovine serum (FBS), penicillin (100 IU/ml) and streptomycin (100 μg/ml). All cells were grown at 37°C and in the presence of 5% CO2.

We purchased TNF-α (R&D systems), 3-deazaneplanocin A (DZNep, Cayman), α-CD3/CD28 antibodies conjugated to magnetic beads (Dynal Biotech) and IL-2 (R&D Systems, Inc.).

### HIV lentiviral vectors and generation of VSV-G-pseudotyped viral particles

The HIV-1 based lentiviral vectors pHR′P-PNL-mCherry and pHR′P-PNL-d2EGFP were constructed with either with wild-type Tat or defective H13L Tat as described previously (Tyagi and Karn, 2007). The construction of pHR′P-Luc has also been previously detailed (Tyagi and Karn, 2007). The shRNA vector pHR′P-SIN-CMV-GFP was generated by cloning the CMV-GFP insert into the *Sac*II to *Xho*I sites of the pHR′P-SIN-18 vector. The short hairpin RNA (shRNA) inserts were initially cloned into the pSuper vector (Oligoengine). The shRNA plus the H1 promoter were then cloned into pHR′P-SIN-CMV-GFP between the *Bam*H1 and *Sal*I sites as earlier detailed (Tyagi and Karn, 2007; Zufferey et al., 1998). The HIV-based lentiviral vector particles pseudotyped with the vesicular stomatitis virus glycoprotein (VSV-G) were produced using a three-plasmid co-transfection procedure (Dull et al., 1998; Naldini et al., 1996). The viruses were concentrated by ultracentrifugation, aliquoted and frozen at −80°C until use.

### Generation of latently infected primary CD4^+^ T cells

The latently infected primary CD4+ T cells were generated using our well established methodology (Tyagi et al., 2010). Briefly, the purified CD4^+^ T cells (>98% pure) either from PBMCs or tonsils were stimulated for 4 days with 25 μl of α-CD3/CD28 antibodies conjugated to magnetic beads (Dynal Biotech) per million cells along with 20 U/ml of IL-2. One million cells were infected with one of the VSV-G-pseudotyped HIV viruses. After 2 to 4 days, the fluorescent cells were purified by fluorescence-activated cell sorting (FACS). The pure population of infected cells was again amplified in the presence of α-CD3/CD28 antibody-conjugated Dynal beads (25 μl/10^6^ cells) and 20 U/ml of IL-2 for 2 to 3 weeks. Fresh medium was added every 4 days to maintain a density of 1.5 × 10^6^ to 2.0 × 10^6^ cells/ml. Once become 0.5 × 10^8^ to 1 × 10^8^, the cells were placed on 30 to 40% confluent H80 adherent cell mono-layer (Sahu et al., 2006; Tyagi et al., 2010). Every 2 to 3 days, half of the culture medium was replaced by fresh IL-2-containing medium, and every 2 weeks the T lymphocytes were transferred to the fresh flasks of H80 feeder cells.

### ChIP assays and q-PCR

Chromatin immunoprecipitation (ChIP) assays were done following a previously described protocol (Tyagi and Karn, 2007; Tyagi et al., 2010). To activate cells, we used either 10 ng/ml TNF-α (for cell lines) or 25 μl per 10^6^ cells of α-CD3/CD28 antibodies bound Dynal beads along with 20 U/ml of IL-2 (for primary T cells). The chromatin was immunoprecipitated using different antibodies detailed in the antibodies section. Each sample (5%) was analyzed by quantitative real-time PCR (qPCR) to assess the amount of sample immunoprecipitated by an individual antibody. Control preimmune IgG value was subtracted from each sample value to remove the background counts. SYBR green PCR master mix (12.5 μl/sample; Bio-Rad) combined with 1 μl of each primer, 5 μl of ChIPed DNA and water to a final volume 25 μl was analyzed by real-time q-PCR. The primers used were the following (numbered with respect to the transcription start site): Promoter region of HIV-1 LTR (promoter) forward,−116, AGC TTG CTA CAA GGG ACT TTC C and reverse +4, ACC CAG TAC AGG CAA AAA GCA G; Nucleosome-1 region HIV-1 LTR (Nuc-1) forward +30, CTG GGA GCT CTC TGG CTA ACT A and reverse +134, TTA CCA GAG TCA CAC AAC AGA CG; glyceraldehyde 3-phosphate dehydrogenase (GAPDH) promoter forward, −125, CAC GTA GCT CAG GCC TCA AGA C and reverse, −10, AGG CTG CGG GCT CAA TTT ATA G; GAPDH was also assessed by forward, −145, TAC TAG CGG TTT TAC GGG CG and reverse, +21 TCG AAC AGG AGG AGC AGA GAG CGA.

### Western blotting

Western blotting was performed according to standard protocols. Briefly, nuclear extracts were run on acrylamide Tris-HCl buffered SDS-PAGE gels (7.5% to 10%). Western blots were developed using suitable primary and secondary HRPO-conjugated anti-rabbit or anti-mouse antibodies (Dako, SantaCruz Biotechnology). Primary antibodies are described in the antibodies section.

### Luciferase assays

Cells in six-well plates were harvested after 48 h of treatment, washed twice in phosphate-buffered saline (PBS), and then lysed in 100–200 μl of 1 × Passive Lysis Buffer (Promega) for 30 min at room temperature (RT). The firefly luciferase activity was analyzed by luciferase reporter assay system (Promega) and normalized by protein concentration of cell lysate.

### Transfection

For generating Vesicular stomatitis virus G-pseudotyped HIVs, the 293T cells were transfected with plasmids using Lipofectamine (Invitrogen/ThermoFisher Scientific) applying a previously described methodology (Dull et al., 1998; Naldini et al., 1996). The viral titer was determined by the infection of 1 × 10^6^ Jurkat T cells with serial dilutions of the harvested culture supernatant. However, for the transfection of Small Interfering RNA (siRNA) we used Lipofectamine RNAiMAX (Invitrogen/ThermoFisher Scientific) according to the manufacturer’s protocols. For each gene 4 siRNA target sequences (20 nM each), detailed in the sequences of primers section, were used. As control, cells were either infected with lentiviral vector expressing scrambled shRNA or transfected with a neutral scrambled siRNA sequence. Briefly, the cells were incubated with Lipofectamine RNAiMAX reagent and siRNA in Opti-MEM medium (Invitrogen/ThermoFisher Scientific) at RT for 5 to 7 minutes. Subsequently, siRNA-lipid complex was added to the cells and incubated for 48 hours at 37°C in cell culture incubator. Three independent experiments were performed (error bars = SD; n = 3). The knockdown effects were assessed by Western blotting using respective antibodies.

### Flow cytometry

The expression of fluorescent reporter gene was assessed through fluorescence-activated cell sorting (FACS) using a FACSCalibur flow cytometer. Forty-eight hours post-stimulation via α-CD3/CD28 antibody along with IL-2, the expression of fluorescent protein was assessed. The mixed populations were sorted by flow cytometry to enrich 100% HIV infected cells based on fluorescent protein expression. The shutting down process of latently infected cells to become latent was assessed by flow cytometry every other week (Tyagi et al., 2010). The presence of latent provirus was confirmed by activating the latently infected cells with α-CD3/CD28 antibody along with IL-2 for roughly 50 hours. For some experiments in order to analyze cells afterword, the cells were fixed with 3% formaldehyde and stored at 4°C, before flow cytometry.

### Antibodies

Several of the antibodies were purchased from Santa Cruz, including anti-RNAP II (N-20 sc-899; F-12 sc-55492; A-10 sc-55492), CBF-1(Millipore, AB5790; E-7 sc-271128; Sigma, AB384), CIR (C-19; H-1, sc-514120), mSIN3A (AK-11, G-11 sc-5299), HDAC-1 (H-51 sc-7872; H11 sc-8410, 10E2 sc-81598), HDAC-3 (H-99 sc-11417; A-3 sc-376957), p300 (C-20 sc-585; F-4 sc-48343), HP1α (C15 sc-10210; Active Motif 2HP-1H5).GAPDH (6C5 sc-32233; 0411 sc-47724), anti-β-actin (C-4 sc-47778), Spt5 (D-3 sc-133217), EED (H-300 sc-28701; Active Motif 41D) and p65 (F-6 sc-8008); Preimmune Rabbit IgG control (Cell Signaling, #2729S), anti-phospho-Ser2 RNAP II (Active Motif 3E10; Abcam ab5095), acetyl-histone H3 (Upstate 07-593); SUZ12 (Cell Signaling, 3737S); EZH2 (Cell Signaling 5246S; Millipore 17-662); JARID1A (Abcam, ab65769); H3K9me3 (Upstate 07-442, Abcam ab8898-100); H3K27me3 (07-449; Upstate); RING1B (Active Motif 39663); BMI1(Active Motif AF27, Upstate F6).

### Sequences of siRNAs, shRNA and constructs used

**SUZ12** (GCTGACAATCAAATGAATCAT, GCTTACGTTTACTGGTTTCTT, CCAAACCTCTTGCCACTAGAA, CGAAACTTCATGCTTCATCTA)

**EED** (GACACTCTGGTGGCAATATTT, CCTATAACAATGCAGTGTATA, GTGCGATGGTTAGGCGATTTG, CTGGATCTAGAGGCATAATTA)

**EZH2** (CGGCTCCTCTAACCATGTTTA, CCCAACATAGATGGACCAAAT, GCTGACCATTGGGACAGTAAA, CAACACAAGTCATCCCATTAA)

**BMI1** (ATTGATGCCACAACCATAATA, GGAACCTTTAAAGGATTATTA, CAGCAAGTATTGTCCTATTTG, TAATGGATATTGCCTACATTT)

**Scrambled** (TTGATGCACTTACTAGATTAC)

### CBF-1 shRNA constructs

Besides using Clone ID TRCN0000016204, TRCN0000016203 (Open Biosystems), we also used following shRNA sequences to clone in a lentiviral vector and expressed through H1 promoter (Tyagi and Karn, 2007).

1.CCGGCCAGGATAACTGTGCGAACATCTCGAGATGTTCGCACAGTTATCCTGGTTTTTG

2.CCGGGCTGGAATACAAGTTGAACAACTCGAGTTGTTCAACTTGTATTCCAGCTTTTT

### Statistical analysis

Data were analyzed using Microsoft Excel or GraphPad Prism softwares. For paired samples, statistical analyses were performed using Student’s *t* test. One-way analysis of variance (ANOVA) was performed for multiple data point comparisons.

## Results

### CBF-1 knockdown disrupts the latency maintenance and leads to the proviral reactivation in primary T cells

We have already confirmed the important role of CBF-1 during the establishing phase of HIV latency, including in primary CD4+ T cells (Tyagi and Karn, 2007; Tyagi et al., 2010). There is an inverse correlation between cellular levels of CBF-1 and HIV gene expression. Accordingly, cells harboring latent provirus have higher level of CBF-1. However, upon cell activation, we observed a sharp decline in the cellular levels of CBF-1 and a corresponding reactivation of latent provirus. Moreover, using Jurkat cells, a T cell line, the important role of CBF-1 during the maintenance phase of HIV latency was illustrated (Tyagi and Karn, 2007). In order to extend those studies and further define the role of CBF-1 in primary CD4+ T cells during latency maintenance, we performed some knockdown experiments. We knockdown the endogenous CBF-1 in physiologically relevant primary CD4+ T cells carrying latent provirus and reactivation of latent provirus was assessed. The rationale behind doing these experiments was if CBF-1 imposed transcriptional restrictions plays role in HIV latency maintenance, then its removal should relieve those restrictions and lead to proviral reactivation.

The latently infected primary CD4+ T cells harboring pHR’P-Luc HIV provirus, which expresses luciferase as reporter through LTR promoter (Fig. 1A), were generated using established methodology (Tyagi and Karn, 2007; Tyagi et al., 2010). To knockdown endogenous CBF-1, the latently infected primary cells were superinfected with lentiviral vectors expressing shRNAs either against CBF-1 or control scrambled shRNA. Scrambled shRNA was confirmed for its neutrality towards HIV and cellular genomes using the NCBI program Blast (Tyagi and Karn, 2007). More than 70% knockdown was obtained using 4 μg of shRNA vectors (Fig. 1C). Depletion of CBF-1resulted in reactivation of latent provirus, indicated by the enhanced expression (more than three-fold) of luciferase reporter gene compared to a scrambled shRNA control (Fig. 1B). As positive control to show the population of cells carrying recativable latent provirus in their genome, cells were activated through T cell receptor (TCR) stimulation by treating the cells with anti-CD3/-CD28 antibodies in the presence of IL-2 (α-CD3/-CD28/IL-2). Following TCR stimulation, we noted more than twice luciferase counts than those obtained upon CBF-1 knockdown. Notably, as detailed above we also observed reduction to cellular levels of CBF-1 following cell activation via TCR stimulation (Fig. 1C).

**Figure 1:**
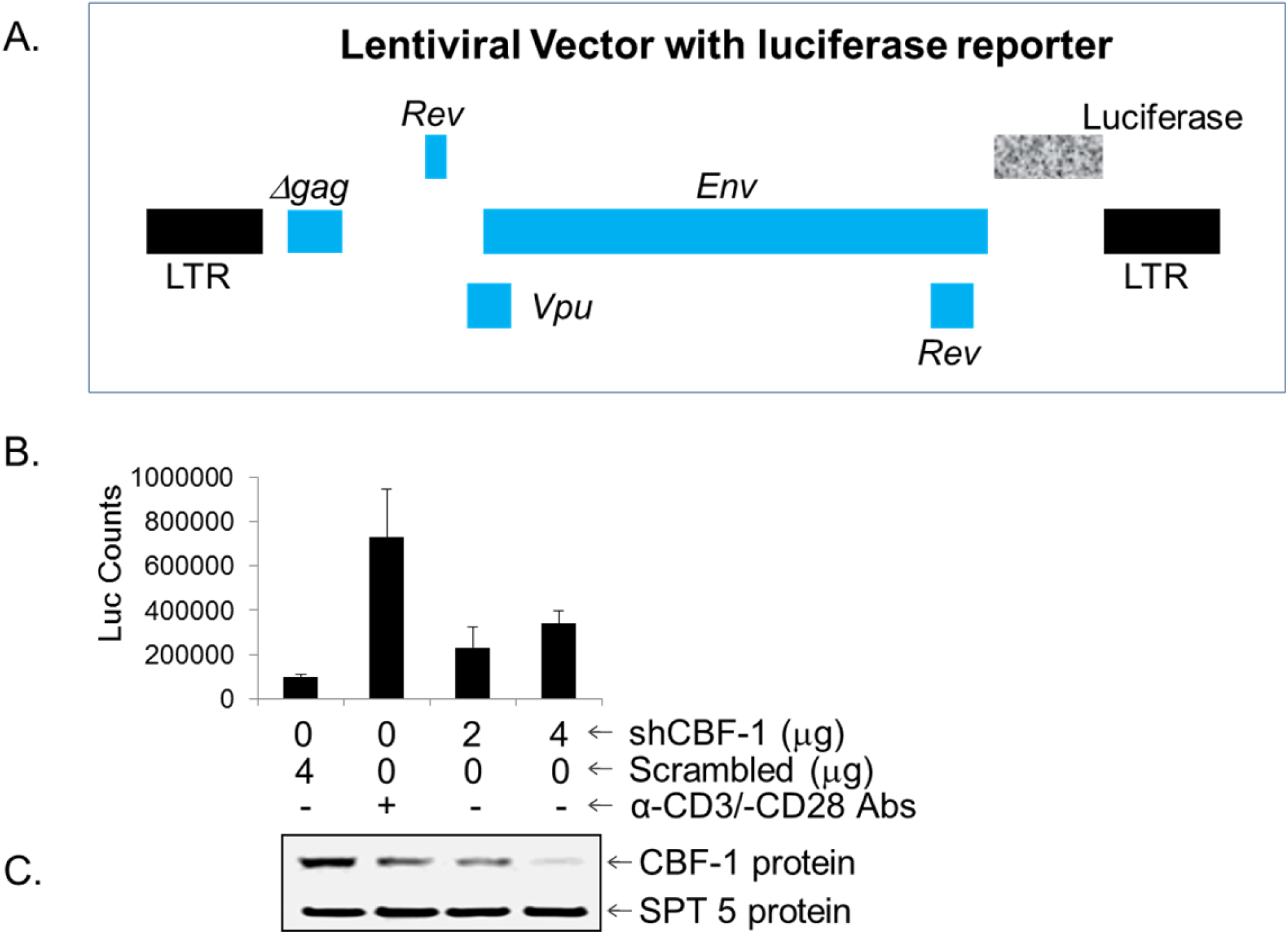
Knockdown of CBF-1 in primary CD4+ T cells reactivates latent HIV proviruses. (**A**) Structure of lentiviral vector (pHR’P-Luc) which carries reporter *luciferase* gene under HIV LTR promoter. (**C**) Western Blot demonstrating CBF-1 knockdown in cells expressing shRNAs against CBF-1 and control cells expressing scrambled shRNA. (**B**) Luciferase assay showing proviral reactivation in cells with pHR’P-Luc that are superinfected with different amounts of lentiviral vectors expressing either shRNAs against CBF-1 or control scrambled shRNA. Error bars represent the SEM of three separate experiments.

Partial reactivation of latent provirus after CBF-1 depletion indicated the involvement of additional factors in restricting HIV gene expression during the maintenance phase of HIV latency. Moreover, besides epigenetic restrictions other mechanisms also play role in restricting HIV in the latent state (Hakre et al., 2011; Mbonye and Karn, 2014; Taube and Peterlin, 2013).

These results in primary T cells along with our previously published data obtained using T cell lines (Tyagi and Karn, 2007) verified the important role CBF-1 during the maintenance phase of HIV latency. Hence, CBF-1 besides inducing the establishment of HIV latency (Tyagi and Karn, 2007), promotes the maintenance of HIV latency.

### CBF-1 promotes HIV latency by inducing multiple types of repressive epigenetic modifications at HIV LTR

In earlier publications we demonstrated that CBF-1 restricts HIV transcription by recruiting HADCs containing corepressor complexes at HIV LTR. HDACs subsequently mediate the deacetylation of core histones, which eventually facilitates the establishment of HIV latency, both in transformed and primary CD4+ T cells (Tyagi and Karn, 2007; Tyagi et al., 2010). Since various corepressor complexes contain HDACs, a goal was to define the precise identity of the corepressor complex recruited by CBF-1 at HIV LTR during HIV latency. In addition to histone deacetylation, numerous studies including ours have established the importance of other repressive epigenetic modifications, such as tri-methylation of core histone H3 at position 9 (H3K9me3) and 27 (H3K27me3) during HIV latency (du Chene et al., 2007; Imai et al., 2010; Mbonye and Karn, 2011; Tyagi and Bukrinsky, 2012; Tyagi and Karn, 2007; Tyagi et al., 2010). The role of enzymes responsible for catalyzing these epigenetic modifications during HIV latency has also been well documented (du Chene et al., 2007; Friedman et al., 2011; Imai et al., 2010; Marban et al., 2005; Marban et al., 2007; Pearson et al., 2008). We have also confirmed the role of H3K9me3 and H3K27me3 in primary CD4+ T cells, during HIV latency (Tyagi et al., 2010). We investigated if CBF-1 recruited corepressors are responsible for inducing those repressive histone H3 methylations and promoting HIV latency.

In order to determine whether CBF-1 is responsible for inducing varying repressive epigenetic modifications, we assessed the impact of CBF-1 knockdown on the resultant epigenetic changes at HIV LTR. If the enzymes present in CBF-1 recruited corepressor complex are responsible for catalyzing H3K9me3 and H3K27me3 modifications, then CBF-1 knockdown should result in reduced recruitment of the corepressor complex and thus less generation of H3K9me3 and H3K27me3 at HIV LTR.

The latently infected Jurkat T cell line, clone E4, in which short lived green fluorescent protein (d2EGFP) reporter is replaced with the Nef gene, was used (Pearson et al., 2008; Tyagi and Karn, 2007). The latently infected cells were superinfected with lentiviral vectors carrying shRNAs either against CBF-1 or control scrambled shRNA, a target neutral sequence. The cellular population carrying shRNA vectors was enriched via puromycin selection.

To assess the binding of different transcription and epigenetic factors, besides changes in corresponding epigenetic modifications at HIV LTR before and after CBF-1 knockdown quantitative ChIP assays were performed and the two critical regions of HIV LTR, promoter (Fig. 2A) and nucleosome-1 (Nuc-1) (Fig. 2B) were examined. The immunoprecipitated DNA was measured through q-PCR using primer sets directed to the promoter region (-116 to +4) and nucleosome-1 region (+30 to +134) of LTR with respect to transcription start site. The Immunoprecipitant of control IgG was subtracted from all samples as background. To provide a control for equal loading, the results were normalized with housekeeping GAPDH gene expression (-145 to +21), a constitutively expressing cellular gene. Latently infected Jurkat cells showed low levels of RNAP II at both the promoter and Nuc-1 regions of LTR (Fig. 2). Given ChIP resolution capacity of ∼500 bp due to DNA shearing limit during sonication, we found overlapping signal at the promoter and neighboring Nuc-1 regions of LTR. Nevertheless, histone modifications were clearly more prevalent at the histone rich Nuc-1 region (Fig. 2B). The lower amount of RNAP II at LTR of latent provirus confirms that latent HIV proviruses are restricted in HIV transcription. As anticipated, we found higher levels of CBF-1 and HDAC-1 at latent provirus in accord with our previous studies that demonstrate that after binding to LTR, CBF-1 recruits HDACs containing corepressor complexes (Tyagi and Karn, 2007). We also observed the accumulation of other heterochromatic marks, H3K9me3 and H3K27me3 at HIV LTR of latent provirus. This observation verifies that CBF-1 promotes transcriptional silencing of latent provirus by inducing multiple layers of repressive epigenetic modifications at HIV LTR.

**Figure 2:**
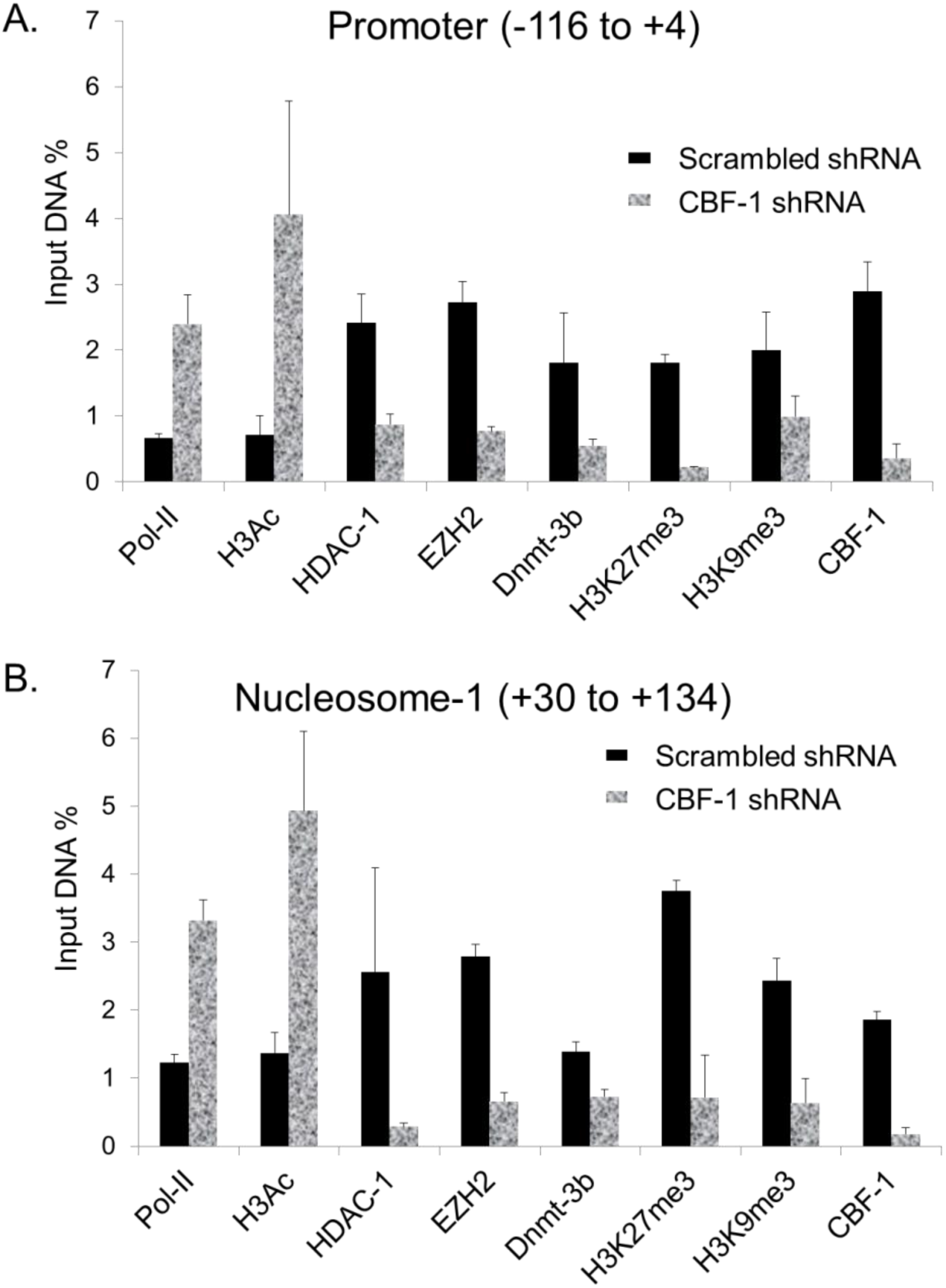
CBF-1 restricts HIV transcription by inducing multiple types of repressive epigenetic modifications at HIV LTR. ChIP analyses were performed using latently infected Jurkat T cells to evaluate the turnover of different epigenetic modifications at HIV LTR before and after knockdown of endogenous CBF-1 using the indicated antibodies. Primer sets directed to the (**A**) Promoter region (-116 to +4) with respect to transcription start site; **(B)** Nucleosome 1 (+30 to +134) with respect to transcription start site of HIV-1 LTR. Black bars: latently infected cells; Shaded gray bars: CBF-1 knockdown cells. The depicted ChIP assay results were reproduced 5 times.

Interestingly, following CBF-1 knockdown the level of CBF-1 at HIV LTR drops sharply, confirming that there is less amount of CBF-1 in the cell for recruitment to LTR. As anticipated, we found parallel dissociation of HDACs containing corepressor complexes from LTR, demonstrated by the removal of HDAC-1, further illustrating the direct role of CBF-1 in recruiting HDACs. The loss of HDACs resulted in enhanced acetylation of core histones, represented by the hyperacetylation of core histone H3 following CBF-1 knockdown. Notably, there is corresponding loss of other repressive epigenetic modifications from LTR following CBF-1 knockdown, namely H3K9me3 and H3K27me3. This indicates that the enzymes present in the CBF-1 recruited corepressor complex are responsible for catalyzing these epigenetic modifications. In fact, following CBF-1 knockdown we found the corresponding loss of EZH2 from LTR, an enzyme that catalyzes H3K27me3 and the core component of PRC2 corepressor complex. On the other hand, the establishment of the epigenetic mark H3K9me3 is usually catalyzed by SUV39H1 and G9A (Kouzarides, 2002, 2007). The presence of SUV39H1 and G9A at HIV LTR and their role during HIV latency have been well documented (du Chene et al., 2007; Imai et al., 2010; Marban et al., 2007; Nguyen et al., 2017). Both SUV39H1 and G9A are known to interact with various subunits of PRC1 (Li et al., 2010; Sewalt et al., 2002), suggesting the presence of PRC1 as well. Together, these findings suggested that along with PRC2, CBF-1 brings PRC1 to HIV LTR during HIV latency.

### CBF-1 promotes both the establishment and the maintenance of HIV latency by recruiting PRCs at HIV LTR

The presence of PRC2 at HIV LTR and its role during HIV latency establishment and maintenance has been well documented (Friedman et al., 2011; Matsuda et al., 2015; Nguyen et al., 2017; Tripathy et al., 2015). However, factors involved in the recruitment of PRCs at LTR are not fully defined. JARID2 has been shown to facilitate the recruitment of PRC2 at numerous cellular genes (Margueron and Reinberg, 2011; Pasini et al., 2010). Similarly, Karn’s laboratory has shown that JARID2 also promotes the recruitment of PRC2 at HIV LTR (Nguyen et al., 2017). Thus, suggesting that several factors can play role in recruiting PRCs at HIV LTR. We evaluated the recruitment profile of the main core components of PRC1 and PRC2 corepressor complexes at HIV LTR, before and after knocking down the endogenous CBF-1 protein. The levels of different factors were assessed by performing quantitative ChIP assays (Fig. 3). The latently infected Jurkat cells were transduced with lentiviruses expressing shRNAs either against CBF-1 or neutral scrambled shRNA. We determined the recruitment kinetics of different core components belonging to PRC1 and PRC2 corepressor complexes at two critical regions of HIV LTR, promoter and Nuc-1. As indicated in Figure 3, we detected higher levels of core components of PRC1 and PRC2 complexes at the LTR of latent HIV provirus, namely EED, SUZ12, EZH2 and BMI1. The detection of EED, SUZ12, and EZH2 marks the presence of PRC2 while the recruitment of BMI1 indicates the presence of PRC1. This result shows that latent provirus accumulates both PRCs at its LTR, and suggests the role of PRCs during HIV latency. However, upon CBF-1 knockdown, we observed the corresponding dissociation of core components of both PRCs complexes from LTR (Figs. 3A and 3B). The parallel loss of core components of PRC1 and PRC2 following CBF-1 knockdown confirms that CBF-1 is responsible for their recruitment at HIV LTR. Thus, our results convincingly demonstrate that CBF-1 promotes HIV latency by recruiting PRC1 and PRC2 at HIV LTR.

**Figure 3:**
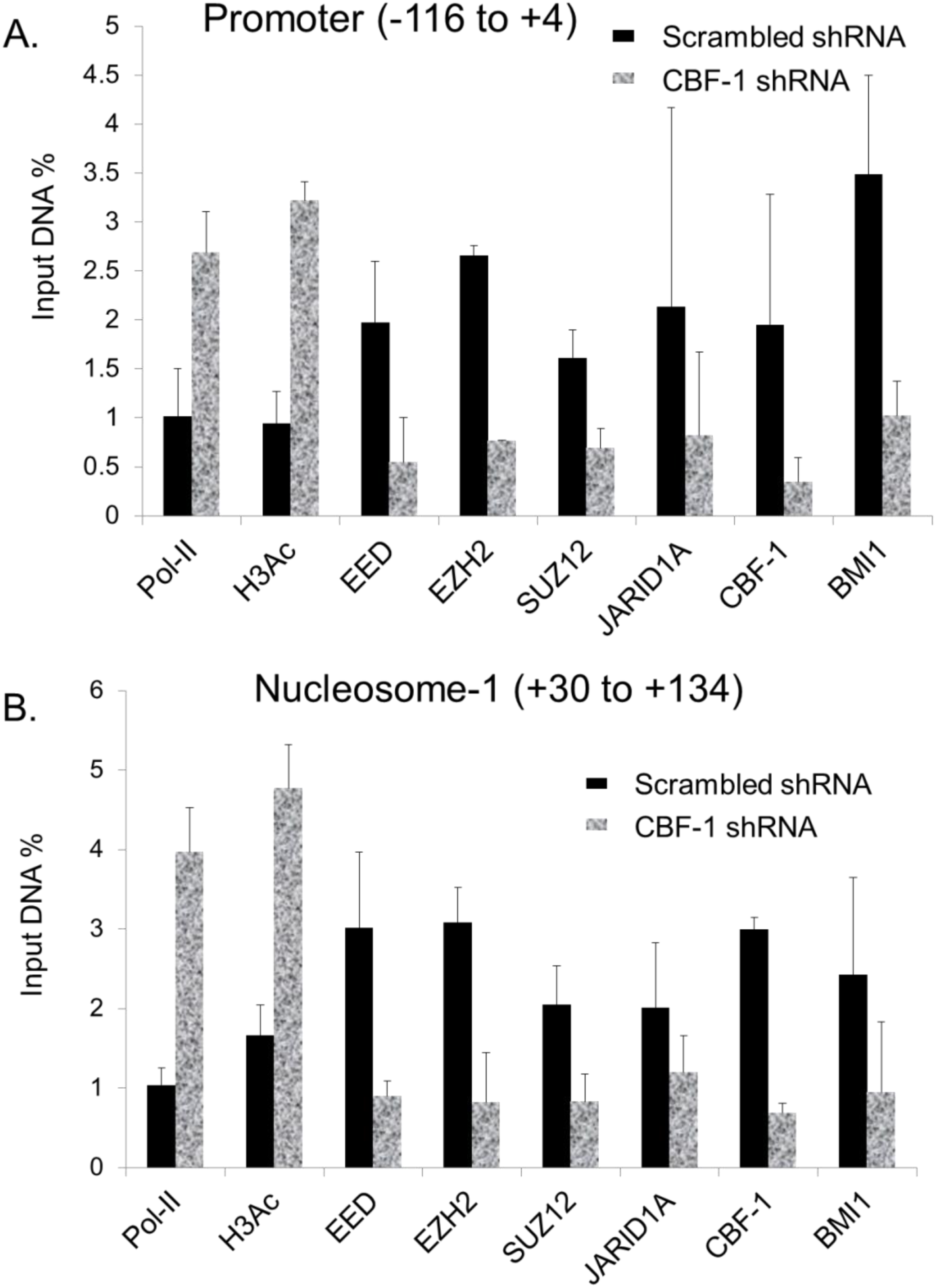
CBF-1 knockdown resulted in dissociation of different factors belonging to both PRCs (PRC1 and PRC2). ChIP analyses were performed using latently infected Jurkat T cells before and after knocking down the endogenous CBF-1. CBF-1 knockdown leads to the dissociation of various core components of both PRCs showing the role of CBF-1 in their recruitment at HIV LTR. (**A**) Promoter region (-116 to +4); (**B**) Nucleosome 1 (+30 to +134). Black bars: latently infected cells; Shaded gray bars: CBF-1 knockdown cells. Error bars represent the SEM of three independent experiments and three separate qPCR measurements from each experiment.

We also found the presence of JARID1A at HIV LTR. JARID1A is a histone H3K4me3 demethylase, which removes the methyl group from histone H3 at position 4 (H3K4me3), a euchromatic mark. At cellular promoters, JARID1A has been shown to interact with both CBF-1 and recruited corepressor complexes, including PRCs (Liefke et al., 2010; Pasini et al., 2008; van Oevelen et al., 2008). Accumulation of epigenetic modifications, such as H3Ac and H3K4me3 supports the establishment of transcriptionally active or euchromatin structures. We found that JARID1A and HDACs of PRCs remove these (H3K4me3 and H3Ac) pro-euchromatin modifications at HIV LTR. Therefore, PRCs besides harboring the enzymes that induce the formation of transcriptionally repressive heterochromatin structures also carry the enzymes which remove the euchromatin structures, consequently supporting prolonged or permanent gene silencing. Hence, by recruiting PRCs at HIV LTR, CBF-1 not only facilitates the establishment of HIV latency, but also promotes the maintenance or stabilization of HIV latency. This conclusion is also supported by the observation that when we knockdown the endogenous CBF-1, the latent provirus gets reactivated (Fig. 1).

Thus, by recruiting PRCs at HIV LTR, CBF-1 induces multiple layers of repressive epigenetic modifications to form transcriptionally repressive heterochromatin structures at HIV LTR during HIV latency. Consequently, CBF-1 not only promotes, but also stabilizes the silencing of latent proviruses.

### CBF-1 recruited PRCs facilitate HIV latency in primary CD4+ T cells

Like an ideal repressor of HIV transcription, CBF-1 is present in abundant amounts in resting T cells, however, upon cell activation CBF-1 levels drops sharply, a property also visible in figure 1C. This unique characteristic of CBF-1 has been confirmed in different cell types (Tyagi and Karn, 2007). This implies that after cell activation, reduced cellular levels of CBF-1 result in poor recruitment of CBF-1 at LTR. In parallel, if CBF-1 contributes to the recruitment of PRCs at LTR, then we envisioned proportionally reduced recruitment of PRCs at LTR. Thus, to validate our hypothesis and provide physiological relevance to these findings, we performed ChIP assays using latently infected primary CD4+ T cells (Figs. 4B to 4G). The latently infected primary CD4+ T cells, harboring either pHR’-PNL-H13LTat-mCherry (Figs. 4B and 4C), pHR’-PNL-wildTat-mCherry (Fig. 4D and 4E) or pHR’-PNL-H13LTat-d2GFP (Figs. 4F and 4G) were generated as described earlier (Pearson et al., 2008; Tyagi and Karn, 2007; Tyagi et al., 2010). These HIV-derived vectors (Fig. 4A) express fluorescent protein reporter genes (either the short-lived d2EGFP or mCherry) in place of the *nef* gene, as detailed earlier (Pearson et al., 2008; Tyagi and Karn, 2007; Tyagi et al., 2010). These viruses express the regulatory proteins Tat and Rev. Thus, like complete HIV the positive feedback circuit that enhances HIV transcriptional elongation and export of mRNA from the nucleus is fully intact. In some of our experiments, in order to increase the frequency of latently infected cells in the population, we utilized Tat carrying the H13L mutation. This partially attenuated Tat variant was originally identified in the U1 latently infected cell line and is highly prevalent in latent proviral pools of HIV patients (Emiliani et al., 1998; Reza et al., 2003; Yukl et al., 2009).

**Figure 4:**
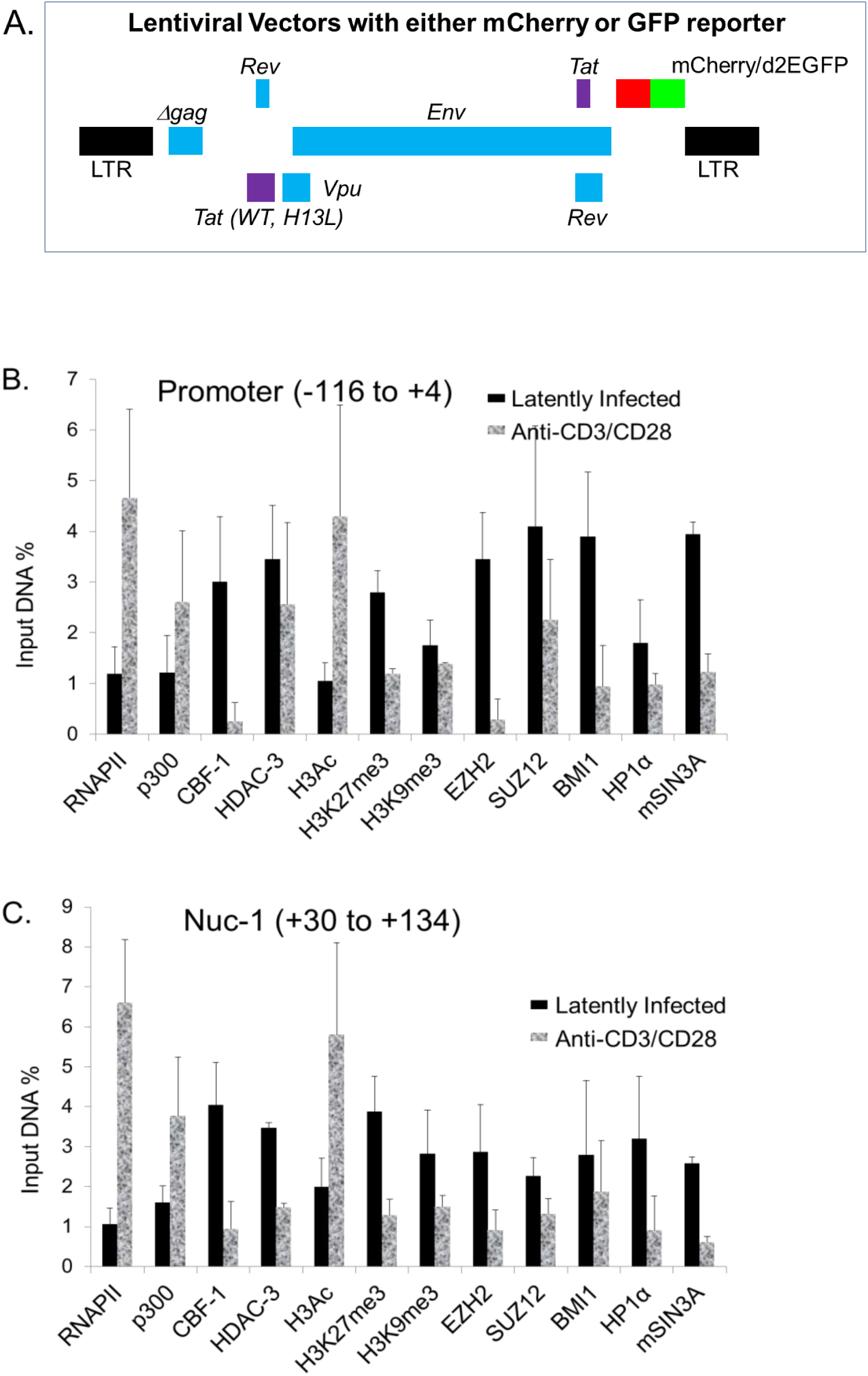
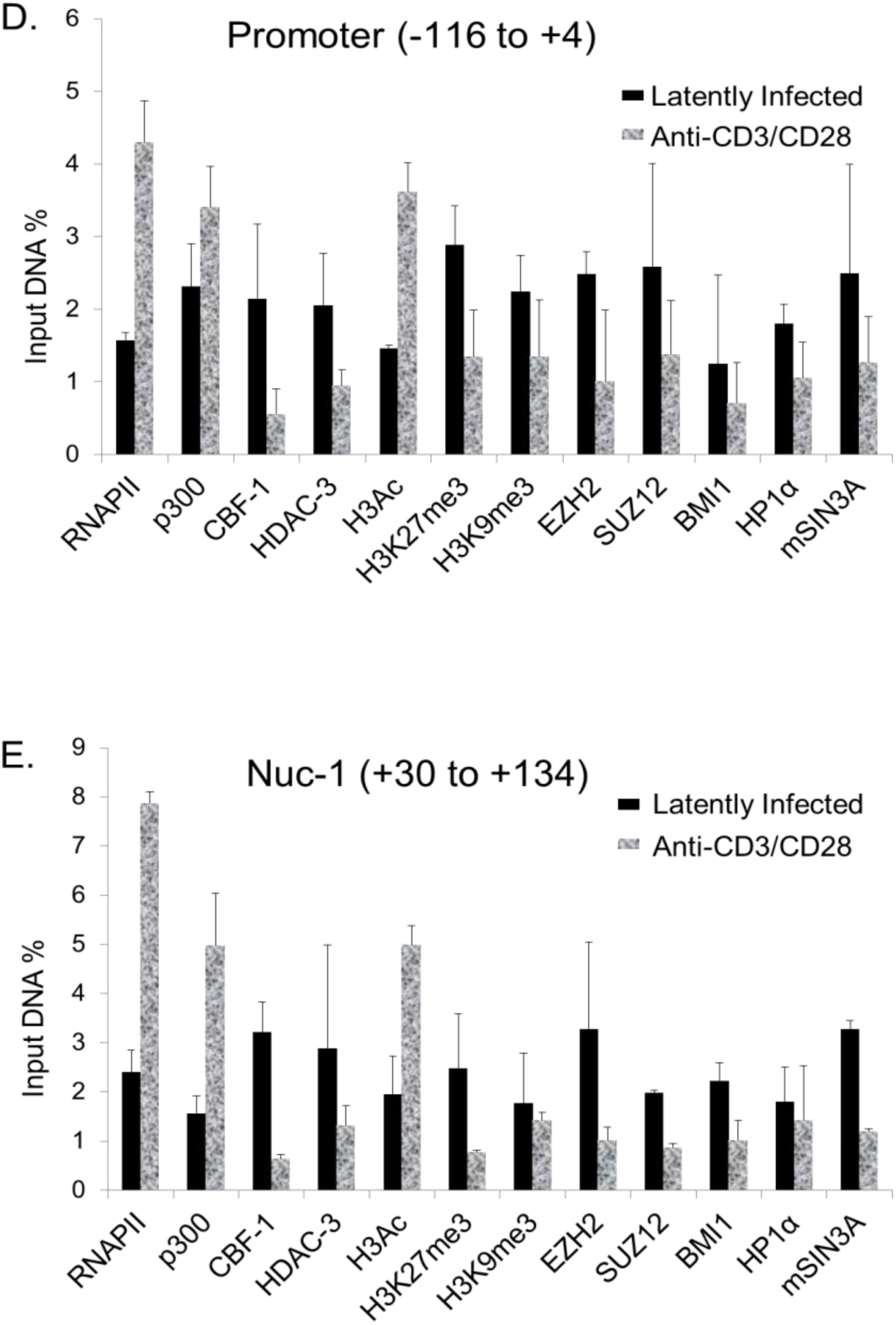
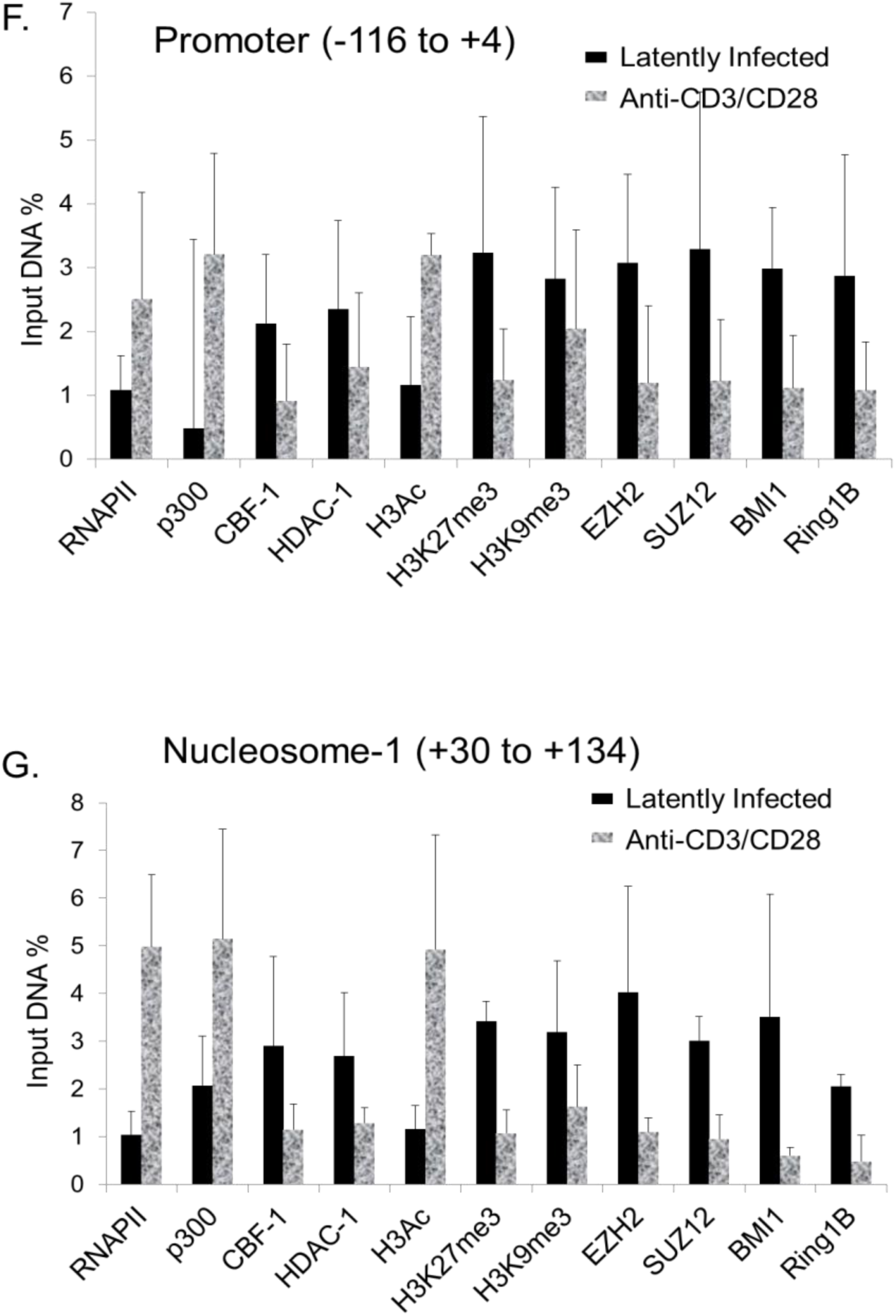
Cell activation leads to the fluctuation in the levels of different chromatin-associated factors that belong to PRC1 and PRC2. ChIP analyses were performed before and after activation of latently infected primary CD4^+^ T cells with α-CD3/CD28 antibodies in the presence of IL-2 for 30 minutes. (**A**) Structure of lentiviral vectors. In some experiments, mCherry was used in place of the d2EGFP fluorescent reporter depicted in this diagram. In order to increase the frequency of latently infected cells in the population, we utilized Tat carrying the H13L mutation, a highly prevalent Tat mutation in latent proviruses. ChIP results were reproduced in three different latency systems harboring different types of proviruses (**B and C**) pHR’-PNL-H13LTat-mCherry, (**D and E**) pHR’-PNL-wildTat-mCherry, (**F and G**) pHR’-PNL-H13LTat-d2GFP. Black bars: latently infected cells; Shaded gray bars: cells stimulated with α-CD3/CD28. Error bars represent the SEM of two independent experiments and three separate qPCR measurements from each analysis.

Quantitative ChIP assays were performed before and after activating the latently infected primary CD4+ T cells with α-CD3/-CD28 antibodies in the presence of IL-2 for 30 minutes. The immunoprecipitated DNA was measured through q-PCR using primer sets directed to the promoter region (-116 to +4) and nucleosome-1 region (+30 to +134) of LTR with respect to the transcription start site. Binding of different transcription factors and epigenetic changes at these regions of LTR dictate overall rate of HIV transcription. The IgG control was subtracted from samples as background. As a control for equal loading in each well, the results were normalized with GAPDH gene expression (-145 to +21), a constitutively expressing cellular gene. As depicted in Figure 4, lower levels of RNAP II were present at the promoter and Nuc-1 regions of latent provirus, validating highly restricted gene expression from LTR promoter of latent provirus. However, in the case of cells infected with provirus carrying wild-type Tat, we observed comparatively higher levels of RNAP II (Figs. 4D and 4E). The reason behind this anomaly is that this cell population consists of around 70% latently infected cells (Figs. 4D and 4E), compared to around 95% latently population in the case of cells infected with provirus carrying H13L Tat (Figs. 4B, 4C, 4F and 4G). The overall LTR binding profiles of different factors were quite comparable in case of cells harboring latent provirus either with wild-type or H13L Tat. We found higher levels of CBF-1, its binding partner mSIN3A and HDAC-1 and -3 at latent proviral LTR in primary T cells. In parallel we found that the LTR of latent HIV contains stable heterochromatin structures, indicated by the higher presence of Histone H3 deacetylation, H3K9me3 and H3K27me3. Analogous to transformed T cells, we found higher recruitment of both PRC1 (BMI1 and RING1) and PRC2 (EZH2 and SUZ12) at the latent HIV LTR (Fig. 4). These findings validate the vital role of PRCs during HIV latency in primary T cells. Reduced levels of RNAP II marks restricted ongoing HIV transcription from LTR. However, TCR stimulation, a condition that results in reactivation of latent provirus (Tyagi et al., 2010), led to a five to seven-folds increase in RNAP II levels at both promoter and Nuc-1 regions. Higher recruitment of RNAP II marks enhanced ongoing HIV transcription after TCR stimulation. Concomitantly, we also observed substantial loss of CBF-1 and recruited corepressor complex, PRCs, from LTR indicated by the loss of mSIN3A, HDAC-1, HDAC-3, EZH2, SUZ12, BMI1 and RING1B (Fig. 4). Loss of HDACs from LTR translated into enhanced acetylation of core histones, indicated by four to seven-fold increase in the acetylation of histone H3 present at the LTR. Similar to our previous observations in primary T cells (Tyagi et al., 2010), we found removal of repressive H3K9me3 and H3K27me3 heterochromatic marks from LTR, recruitment of histone acetyltransferase, p300 at HIV LTR following TCR stimulation. As noted earlier in primary T cells (Tyagi et al., 2010), we found enhanced recruitment of NF-κB (p65) at the promoter region of LTR, as NF-κB binding sites reside in that region (data not shown).

Given the ChIP resolution limit of ∼500 bp, an overlap of signals between adjacent regions, such as promoter and Nuc-1, was expected. Therefore, to some extent we observed similar histone changes at both LTR regions. Nevertheless, notable difference in the levels of histone modifications was clearly visible in Nuc-1 region of the proviruses. These results are consistent with previous studies using transformed cell lines, which have shown that the HIV promoter region is relatively devoid of histones (Tyagi and Bukrinsky, 2012; Tyagi and Karn, 2007; Tyagi et al., 2010; Verdin et al., 1990; Verdin et al., 1993).

In summary, these results have shown that CBF-1 restricts HIV transcription by recruiting both PRC1 and PRC2 during HIV latency in primary CD4+ T cells. We thus validated the role of CBF-1 and PRCs during both the establishment and the maintenance of HIV latency in primary CD4+ T lymphocytes.

### PRCs play direct role in sustaining HIV provirus in latent state

In order to establish the direct role of the PRCs in controlling HIV latency, we knocked down some of the core components of both PRC1 and PRC2. Subsequently, we examined if the removal of repression posed by PRCs on HIV transcription leads to the reactivation of the latent HIV proviruses. Jurkat cells harboring latent HIV provirus pHR’-P-Luc, a lentiviral vector carrying the luciferase reporter gene under the control of the HIV LTR, were used. The cells were transfected with the 20nM siRNA against main components of PRCs (PRC1 and PRC2). We used a mixture of 4 siRNAs (20nM each) against each target gene, and the non-targeting scrambled siRNA was used as control. Efficient knockdown (more than 70%) was quite evident when compared with scrambled control (Fig. 5B). The reactivation of latent provirus was assessed through luciferase reporter assay. Around three-fold reactivation of latent provirus was observed following knockdown of each component of PRCs (Fig. 5A). Notably, the knockdown of the components of PRC2 was slightly more effective than PRC1 subunits in reactivating latent proviruses. Quite similar effects in each case suggest that the removal of any of the core components of PRCs destabilizes the corepressor complex at LTR.

**Figure 5:**
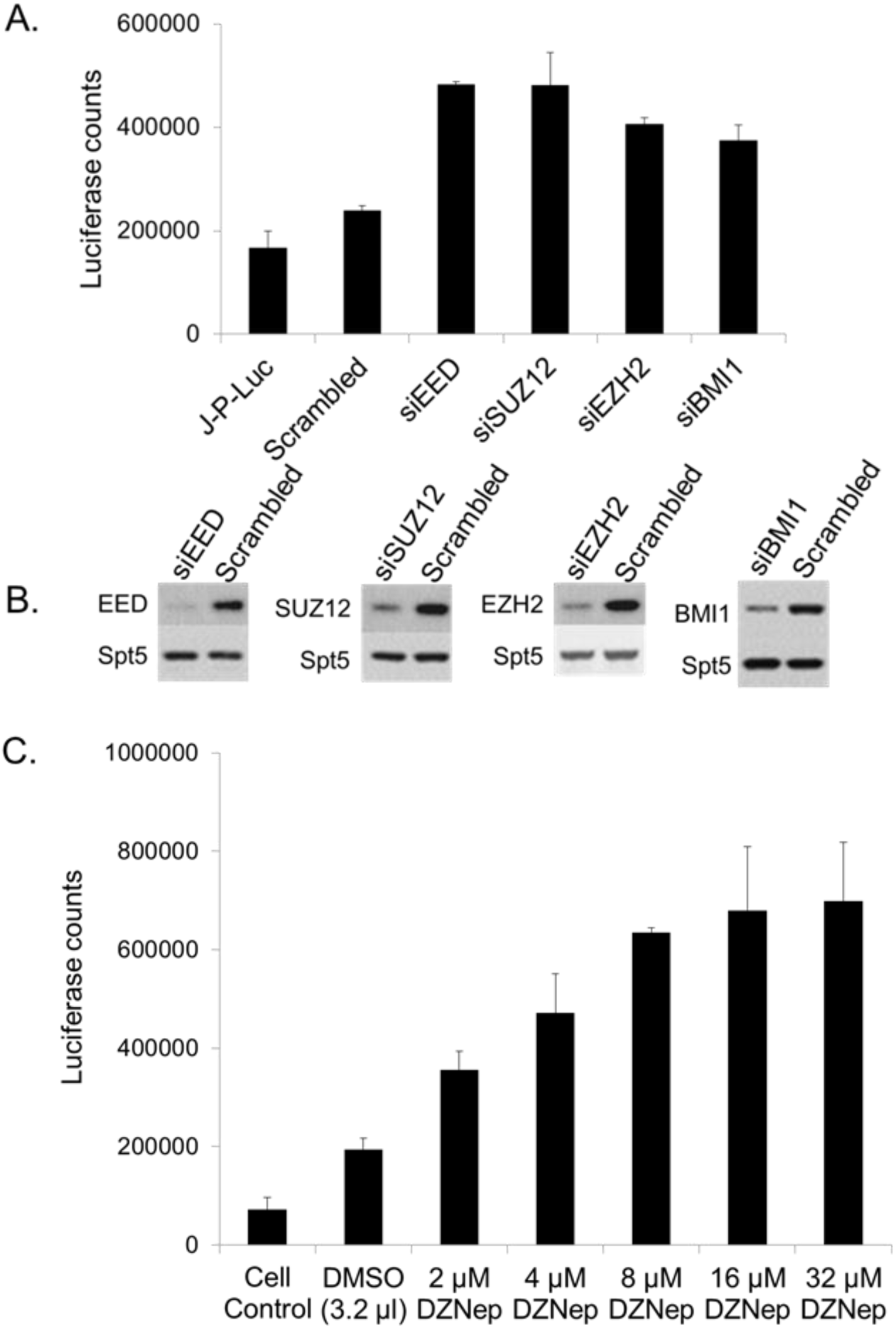
Knockdown of PcG complex led to proviral reactivation. Some of the core PcG complex components were knocked down individually by transfecting latently infected Jurkat-pHR’P-Luc cells with 4 specific siRNAs. The quantification of HIV-1 reactivation of latent provirus was assessed through luciferase assays performed after 52 hours either post siRNA transfection or 48 hours post DZNep treatment. (**B**) Western blot analysis showing the efficiency of siRNA used to knockdown different indicated subunits of PRCs. Quantitative luciferase assays showing proviral reactivation either after knockdown of individual subunits belonging to PRCs (**A**) or upon DZNep treatment (from 2 μM to 32 μM) of cells (**C**). Graphs represent the average and standard deviation from three replicate samples.

Similar results were obtained when we disabled the PRC2 complex by inhibiting the EZH2 using DZNep, a broad-spectrum histone methylation inhibitor. DZNep is known to downregulate the cellular levels of several histone methylases, including EZH2 (Miranda et al., 2009). Latently infected primary CD4+ T cells harboring latent HIV provirus, pHR’-P-Luc, were treated dose-dependently with DZNep (2 μM to 32 μM). After 48 hours cell extracts were assessed for the activity of luciferase enzyme by performing luciferase assays. As anticipated, inhibition of PRC2 by DZNep led to proviral reactivation, which further validates the vital role of PRC2 in promoting HIV latency (Fig. 5C). More than three-fold proviral reactivation was observed at concentrations of 8 μM and beyond. At doses higher than 8 μM more cell toxicity but not much proviral reactivation were observed. These results further corroborated the direct role of PRCs in restricting transcription of latent HIV provirus.

## Discussion

In our previous studies, we demonstrated that CBF-1, after binding to its cognitive sites at HIV LTR, strongly and selectively represses HIV transcription. In this paper, we show that CBF-1 promotes the establishment and maintenance of HIV latency by recruiting Polycomb corepressor complexes at HIV LTR.

The polycomb group (PcG) proteins are divided in the form of two main corepressor complexes, PRC1 and PRC2 (Chittock et al., 2017), that we showed to be present at the HIV LTR. PRCs, by inducing transcriptionally repressive epigenetic modifications, facilitate the assembly of heterochromatin structures at HIV LTR. The PRC1 complex mainly catalyzes the monoubiquitination of histone H2A at lysine 119 residue (H2AK119Ub1) through its Ring subunits, Ring1A/B, which contain E3 ligase activity (Cao et al., 2005; Connelly and Dykhuizen, 2017; Eskeland et al., 2010). On the other hand, PRC2 is primarily characterized by the presence of the histone methyltransferases EZH1/2, which, along with other subunits, mainly SUZ12 and EED, catalyze the di-or trimethylation of histone H3 at lysine 27 (H3K27me2/3) (Cao et al., 2002; Margueron and Reinberg, 2010).

In our previous investigations we have shown the important role of CBF-1 during HIV latency by performing experiments in transformed T cell lines (Tyagi and Karn, 2007). Here, we have extended those findings and confirmed the significant role of CBF-1 during HIV latency in physiologically relevant primary CD4+ T cells. The role of H3K9me3 and H3K27me3 during HIV latency is well established. Moreover, we have shown the importance of these repressive epigenetic marks during HIV latency in primary CD4+ T cells (Tyagi et al., 2010). The formation of the epigenetic mark H3K27me3 is mainly catalyzed by EZH2 enzyme. We noted higher levels of EZH2 and deposition of H3K27me3 at the LTR of latent provirus present in primary T cells. EZH2 is the core component of PRC2 and the Karn’s group has convincingly demonstrated the presence and role of EZH2 and of PRC2 during HIV latency establishment and maintenance (Friedman et al., 2011; Nguyen et al., 2017). These results have been validated by other groups (Matsuda et al., 2015; Tripathy et al., 2015; Yoon et al., 2014). However, the identity of the factor(s) that recruit PRCs at HIV LTR and promote HIV latency were not well defined. Recently, Karn’s group has demonstrated the role of JARID2 in recruiting PRC2 at HIV LTR (Nguyen et al., 2017). In a similar manner, we have been investigating the role of CBF-1 as a recruiter of PRCs at LTR. We proposed that if CBF-1 is responsible for the recruitment of PRC2 and EZH2, then CBF-1 reduction at LTR should translate to lesser accumulation of PRC2 and of H3K27me3 at HIV LTR. Accordingly, we found the comparable loss of EZH2 and H3K27me3 from LTR upon CBF-1 knockdown (Fig. 2). In later CBF-1 ablation experiments, we observed the corresponding loss of other core components of PRC2, namely EED and SUZ12, from LTR (Figs. 2 and 3). Altogether, these results confirmed the direct role of CBF-1 in recruiting PRC2 at HIV LTR during HIV latency.

Upon CBF-1 knockdown we found a parallel loss of the epigenetic mark H3K9me3, suggesting that CBF-1 recruited corepressor complex also carries the enzyme that catalyzes the H3K9me3 epigenetic modification. Notably, PRC2 does not carry any enzyme that catalyzes H3K9me3, but PRC1 is known to bring SUV39H1 and G9A along with it (Li et al., 2010; Sewalt et al., 2002). SUV39H1 and G9A are the two main enzymes which catalyze the formation of the epigenetic mark H3K9me3 at nucleosomes. This observation suggested that along with PRC2, CBF-1 also brings PRC1 to HIV LTR for the generation of transcriptionally repressive heterochromatin structures at HIV LTR during viral latency. In fact, we observed a comparable loss of BMI1 and RING1B, two core components of PRC1 following CBF-1 knockdown in both T cell line and primary T cells (Figs. 3 and 4). The presence of PRC1 at HIV LTR during latency has also been noted by other investigators (Friedman et al., 2011; Nguyen et al., 2017; Tripathy et al., 2015). However, the factor that brings PRC1 at LTR was not known, until our investigation demonstrated that CBF-1 is the cellular protein, which, after binding to LTR at specific sites, brings both PRC1 and PRC2 to inhibit HIV transcription during HIV latency.

In our earlier investigations, we have shown that in resting T cells, which harbor latent provirus, higher levels of CBF-1 are present. However, upon cell activation, the cellular level of CBF-1 drops sharply and latent HIV proviruses get reactivated (Tyagi and Karn, 2007). Thus, CBF-1 acts as an ideal transcriptional repressor which plays a vital role in regulating HIV latency. Therefore, to further validate that cell activation leads to the decline of cellular CBF-1 levels, we showed lesser recruitment of CBF-1 at LTR and consequently loss of PRCs from LTR upon cell activation. The physiological relevance of these findings is evident since they were reproduced in latently infected primary CD4+ T cells. We validated the presence of PRCs at latent HIV proviruses and confirmed their removal from HIV LTR upon cell stimulation through TCR induction (Fig. 4). Proviral reactivation was indicated by the higher RNAP II recruitment and confirmed through enhanced expression of the reporter gene *luciferase*. Using latently infected primary CD4+ T cells in an earlier study, we have demonstrated the presence of both H3K9me3 and H3K27me3 at the LTR of latent provirus, which drops sharply following TCR stimulation (Tyagi et al., 2010). Similar to our previous findings, we noted the presence of components of both PRCs, representing the presence of PRC1 and PRC2 at latent provirus, which abruptly dissociated from HIV LTR upon TCR stimulation (Figs. 4B to 4G). Notably, besides core components of PRCs, we also found the presence of certain interacting partners or auxiliary factors of PRCs, such as HDACs, mSIN3A and HP1α (Margueron and Reinberg, 2011). These factors either bind directly to PRCs components or to the induced epigenetic modifications, e.g. H3K9me3 modification promotes the recruitment of HP1 proteins.

Subsequently, to confirm the direct role of PRCs during HIV latency, we assessed the reactivation of latent provirus after knockdown of different core components of PRC1 and PRC2 repressor complexes (Fig. 5). If PRCs play a significant role in the silencing of latent HIV provirus, then their removal or reduction by knockdown should relieve that restriction and lead to proviral reactivation. Consistent with this idea, upon knockdown of the core components, PRCs become destabilized and we observed two to three-folds reactivation of latent provirus (Fig. 5). Notably, we observed better proviral reactivation following knockdown of PRC2 components than PRC1, suggesting a primary role of PRC2 components in the stability of corepressor complex recruited by CBF-1. Supporting this observation it has been documented that PRC2/EZH2 induced H3K27 methylation promotes the recruitment of PRC1 to target cellular genes, and the disruption of PRC2 leads to the loss of PRC1 from chromatin targets, but the other way around is not always that effective (Boyer et al., 2006).

Interestingly, upon CBF-1 knockdown, we found the corresponding loss of JARID1A, an enzyme which is known to interact with PRC subunits (Liefke et al., 2010; Pasini et al., 2008). JARID1A is a histone demethylase, which selectively demethylates the histone H3 at position K4, H3K4me3. In contrast to the above mentioned trans-repressive epigenetic changes H3K9me3 and H3K27me3, the trimethylation of histone H3 at lysine 4 (H3K4me3) is a euchromatic mark, an epigenetic modification that promotes the establishment of a transcriptionally active euchromatin structure which support transcription. Thus, JARID1A, by removing H3K4me3, inhibits the generation of the euchromatin structure at LTR. Consequently, certain enzymes such as JARID1A and HDACs that are recruited by PRCs remove euchromatic marks, namely H3K4me3 and acetylation of histones, respectively, to provide stability to gene silencing and restrict transient gene reactivation.

Hence, in addition to promoting the establishment of latency by recruiting PRCs at HIV LTR, CBF-1 promotes the maintenance or stabilization of HIV latency. Moreover, the reactivation of latent provirus following CBF-1 knockdown or TCR stimulation further validates the role of CBF-1 during the maintenance phase of HIV latency. Both CBF-1 knockdown or cell activation reduce cellular CBF-1 levels. Therefore, when we removed the restriction posed by CBF-1 through knocking it down or via TCR stimulation, the latent provirus gets reactivated (Figs. 1 and 4).

For summarizing our results, we propose a model to depict the role of CBF-1 in restricting HIV transcription during latency (Fig. 6). According to our model in the absence of transcription factors such as NF-kB and NFAT in quiescent cells, CBF-1 binds to the specific sites at HIV LTR and recruits PRCs. Enzymes of the PRCs subsequently induce multiple layers of repressive epigenetic modifications and remove transcriptionally active epigenetic modifications. These epigenetic changes subsequently facilitate the generation of transcriptionally repressive heterochromatin structures at HIV LTR. Heterochromatin structures restrict the free flow of transcription factors at HIV LTR, which eventually restrict HIV transcription and stabilize restriction. Thus, CBF-1 facilitates the establishment and maintenance of HIV latency. Following cell activation, the levels of CBF-1 drop, whereas levels of transcription factors, including NF-kB and NFAT, rise in the nucleus displacing CBF-1 and PRCs from LTR. Successively, these factors recruit coactivator complexes at HIV LTR, which then establish the euchromatin environment at HIV LTR facilitating the access of transcription machinery at LTR promoter and thus leading to the reactivation of latent proviruses.

**Figure 6:**
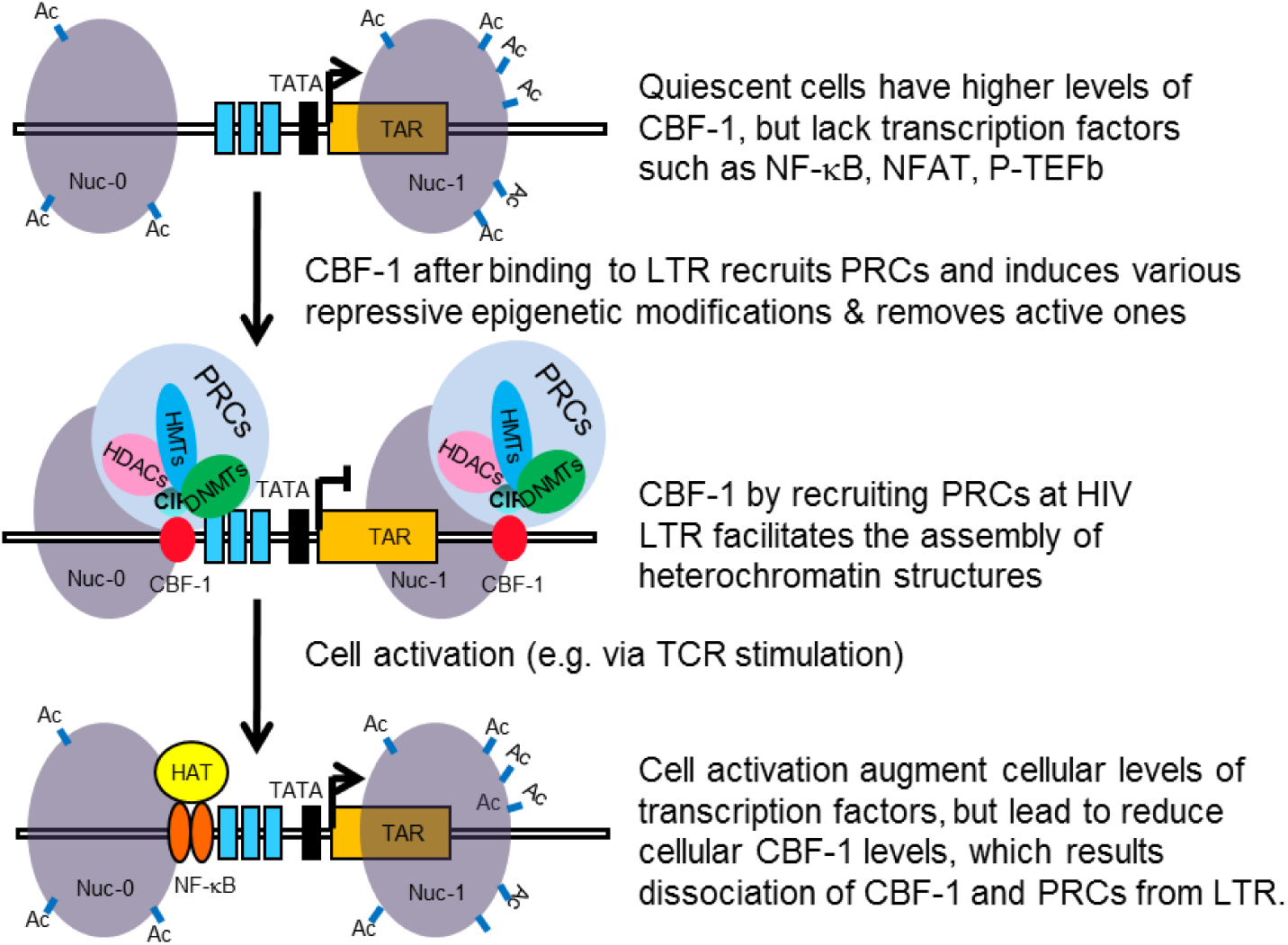
Model of CBF-1 functioning. Based on our findings we propose the following model for the regulation of HIV latency by CBF-1. (**A**) The <u>higher levels of CBF-1 and lack of transcription factors</u> such as NF-kB and NFAT in quiescent cells facilitates the binding of CBF-1 at HIV LTR. (**B**) CBF-1after binding to LTR recruits PRCs, which promote heterochromatin environment at HIV LTR and inhibit the free flow of transcription machinery and thus facilitates the <u>establishment and maintenance of HIV latency</u>. (**C**) Following cellular activation, the levels of CBF-1 drops, but the levels of NF-kB and NFAT rises in nucleus, which displace CBF-1 and corepressor complexes from their binding sites. Subsequently, these factors recruit coactivator complexes at HIV LTR, which then establish the euchromatin environment at HIV LTR that facilitate the access of transcription machinery at LTR promoter and thus lead to the <u>reactivation of latent proviruses</u>.

Taken together our results validated that CBF-1 suppresses HIV gene expression by recruiting both PRC1 and PRC2 at HIV LTR. Hence, we conclude that by recruiting PRCs CBF-1 facilitates both the establishment and maintenance phases of HIV latency. From a clinical standpoint, our findings suggest that for the reactivation of latent proviruses, instead of targeting individual enzymes that induce repressive epigenetic modifications, targeting factors that recruit those enzymes at HIV LTR will result in more profound reactivation of latent provirus. In fact, the removal of the whole corepressor complex will relieve multiple repressive epigenetic modifications simultaneously and could prove to be a better latency reversing strategy, a prerequisite for viral eradication.

## Conclusions

CBF-1 promotes both the establishment and maintenance of HIV latency in primary T cells by recruiting PRC1 and PRC2 at HIV LTR.

## COMPETING INTERESTS

The authors declare that they have no financial competing interests.

## AUTHORS’ CONTRIBUTIONS

Research designed: MT and AC; Research performed: MT, SZ, LSu, KF, AD and LSp; Data analysed: MT, SZ, LSu and LSp; Manuscript written: MT, SZ, AC, AD, GS, LSu, MB and LSp. All authors read and approved the final manuscript.

## ACKNOWLEDGMENTS

We thank the AIDS Research and Reference Reagent Program, Division of AIDS, National Institute of Allergy and Infectious Diseases, US National Institutes of Health; M. Gately (Hoffmann La Roche) for human recombinant interleukin 2; We are also thankful to the Flow Cytometry core facility of George Washington University.

## Funding

The research in Tyagi laboratory is partially funded by National Institute on Drug Abuse (NIDA), NIH Grants, 5R21DA033924-02, 5R03DA033900-02 to MT. This work is also supported by grants of the District of Columbia Center for AIDS Research (DC-CFAR), a NIH-funded program P30AI117970 and startup funds from the George Washington University to MT. The content is solely the responsibility of the authors and does not necessarily represent the official views of National Center for Research Resources or the US National Institutes of Health.

The funders had no role in study design, data collection and analysis, decision to publish, or preparation of the manuscript.

